# Integrative analysis and machine learning based characterization of single circulating tumor cells

**DOI:** 10.1101/867200

**Authors:** Arvind Iyer, Krishan Gupta, Shreya Sharma, Kishore Hari, Yi Fang Lee, Neevan Ramalingam, Yoon Sim Yap, Jay West, Ali Asgar Bhagat, Balaram Vishnu Subramani, Burhanuddin Sabuwala, Tuan Zea Tan, Jean Paul Thiery, Mohit Kumar Jolly, Naveen Ramalingam, Debarka Sengupta

**Affiliations:** Department of Computational Biology, Indraprastha Institute of Information Technology, New Delhi, 110020, India; Department of Computer Science and Engineering, Indraprastha Institute of Information Technology, New Delhi, 110020, India; Centre for BioSystems Science and Engineering, Indian Institute of Science, Bangalore 560012, India; Fluidigm Corporation, 7000 Shoreline Court, Suite 100, South San Francisco, CA 94080, USA; Biolidics Limited, 81 Science Park Drive, 02-03 The Chadwick, Singapore 118257, Singapore; Qualcomm Incorporated, 5775 Morehouse Drive, San Diego, CA 92121, USA; National Cancer Centre Singapore, 11 Hospital Dr, Singapore 169610, Singapore; BioSkryb Corporation, BioLabs, 701 W Main St, Suite 200, Durham, NC 27701, USA; Department of Biomedical Engineering, Faculty of Engineering, National University of Singapore, Engineering Drive 1, Singapore 117575, Singapore; Biomedical Institute for Global Health Research and Technology (BIGHEART), National University of Singapore, 14 Medical Drive, Singapore 117599, Singapore; Center for Artificial Intelligence, Indraprastha Institute of Information Technology, New Delhi, 110020, India; Circle of Life Healthcare Pvt. Ltd., Indraprastha Institute of Information Technology, New Delhi, 110020, India; School of Mathematics, Indian Institute of Science Education and Research, Thiruvananthapuram, 695551, India; Department of Biotechnology, Indian Institute of Technology Madras, Chennai 600036, India; Department of Computational Biology, University of Lausanne (UNIL), Lausanne, 1015, Switzerland; Cancer Science Institute of Singapore, National University of Singapore, Center for Translational Medicine, 117599, Singapore; Guangzhou Institute of Biomedicine and Health, Chinese Academy of Science, Guangzhou, People’s Republic of China

## Abstract

We collated publicly available single-cell expression profiles of circulating tumor cells (CTCs) and showed that CTCs across cancers lie on a near-perfect continuum of epithelial to mesenchymal (EMT) transition. Integrative analysis of CTC transcriptomes also highlighted the inverse gene expression pattern between PD-L1 and MHC, which is implicated in cancer immunotherapy. We used the CTCs expression profiles in tandem with publicly available peripheral blood mononuclear cell (PBMC) transcriptomes to train a classifier that accurately recognizes CTCs of diverse phenotype. Further, we used this classifier to validate circulating breast tumor cells captured using a newly developed microfluidic systems for label-free enrichment of CTCs.

---

A staggering 90% of cancer deaths are attributable to metastases^1^. After detaching from solid tumors, cancer cells travel through the bloodstream to reach distant organs and seed the development of metastatic tumors^2^. Cancer cells under circulation are called circulating tumor cells (CTCs)^3^. As a blood-based bio marker, CTCs offer unabated, real-time insights into tumor evolution and therapeutic responses. Despite these promises, the rareness of CTCs in the peripheral blood hinders their isolation and characterization^3^. Cancers in solid tissues develop from epithelial cells, which are typically densely packed in layers. However, dissemination and migration of cancer cells during metastasis require the acquisition of mesenchymal-like features. Transcendence of epithelial cancer cells into mesenchymal-like ones is popularly known as Epithelial to Mesenchymal Transition (EMT).

It is widely understood that due to the loss of epithelial property only a fraction of CTCs can be expected to express canonical epithelial markers such as Epithelial Cell Adhesion Molecule (EpCAM). The only FDA approved CTC capture platform CELLSEARCH^®^ uses epithelial surface marker EpCAM to detect CTCs in patient blood^4^. Controlled experiments involving cell-lines have shown that recovery of cells with EpCAM expression vary a lot and many canonical epithelial markers are down-regulated in CTCs, undergoing epithelial-mesenchymal transition (EMT)^5^. Therefore, marker-based enrichment techniques are sub-optimal for comprehensive charting of heterogeneous CTC sub-populations. Over the past few years, various CTC capture platforms exploiting biophysical characteristics of cancer cells have been developed^6–8^. CD45-based negative enrichment has also been adopted as an alternative strategy. The potential of such antigen-agnostic platforms have not been fully utilized since the chances of immune cell contamination cannot be completely ruled out^6, 7^. The recent advent of single-cell RNA sequencing (scRNA-seq) has allowed molecular profiling of single CTCs^9^, captured using microfluidic devices^10–14^. Almost all studies that reported molecular profiles of single CTCs resorted to marker based bioinformatic annotation of cell types or applied post-capture staining of CTCs using epithelial/cancer-specific molecular markers^10, 15^. In this study we collated published scRNA-seq datasets of human CTCs and peripheral blood mononuclear cells (PBMCs) to do an integrative analysis and build a machine-learning based classification system that accurately labels CTCs in a marker-agnostic manner and also present the ClearCell^®^ Polaris™ workflow^11, 16^ that involves size-dependant enrichment of CTCs, followed by negative selection based on CD45.

## Results

### Integration of single cell expression datasets of circulating tumor cells

We collected about 700 single CTC transcriptomes from 11 independent studies, representing five different cancer types i.e breast, prostate, lung, pancreas, and melanoma **(Fig 1-b, Supplementary Table-1)**. On the other hand, as control, expression profiles of human PBMCs were collected from six different studies **(Supplementary Table-1)**. About 80% of the CTCs came from various breast cancer studies. CTC datasets that we curated were of variable quality. We preprocessed the data to ensure that the poor-quality cells and unexpressed genes were discarded **(Methods, Supplementary Fig-1)**. We further normalised the combined expression matrix to control for the library depth **(Methods)**. We tracked expression of some of the canonical epithelial and leukocyte markers to cross-validate the cell type identities. Elevated expression levels of a subset of epithelial markers were observed in a vast majority of the CTCs **(Fig 1-c, Supplementary Fig-2)**. Significant up-regulation of platelet and fibroblast markers were observed in large fractions of CTCs **(Fig 1-c, Supplementary Fig-2)**. This combined data source served as the basis for majority of our analysis and development of CTC-immune cell classification system **(Fig 1-a)**.

**Figure 1.**
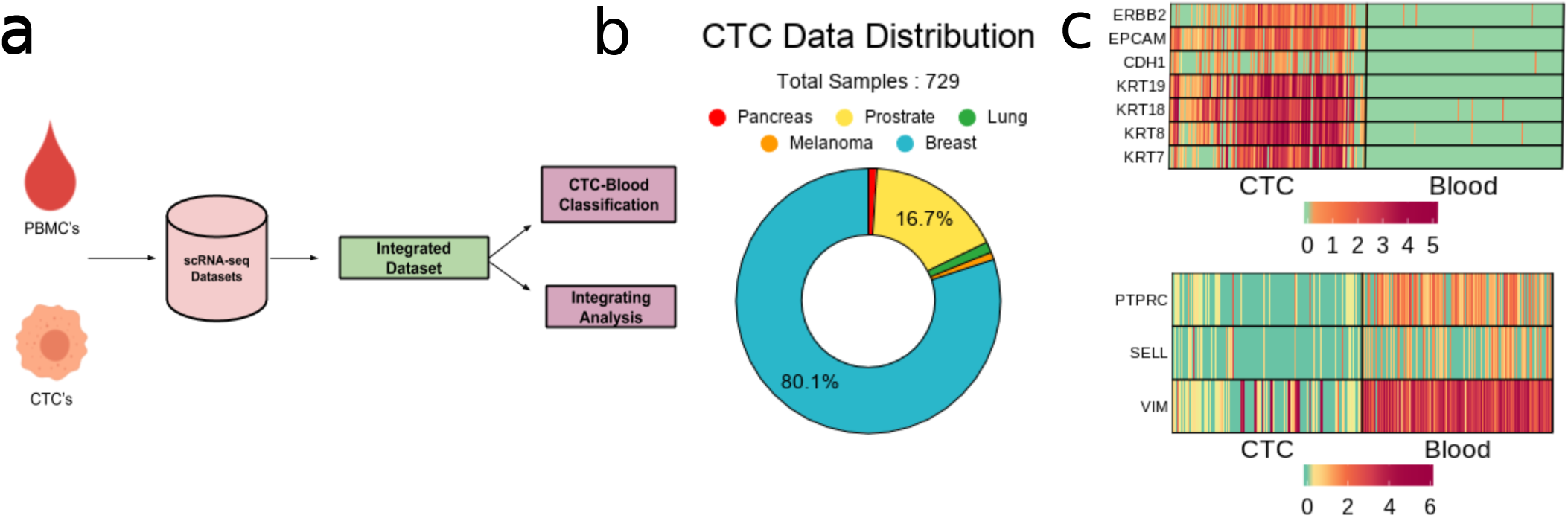
Integrative analysis of CTC transcriptomes: (a) Schematic of study. (b) Cancer types represented by the integrated CTC population. (c) Expression of canonical epithelial and immune cell markers in CTCs and a sub-sample (equal in number as CTCs) of the PBMCs under study.

### Ubiquity of epithelial-mesenchymal transition in cancer metastasis

Epithelial-mesenchymal transition (EMT) and mesenchymal-epithelial transition (MET) have long been postulated to play key roles in cancer metastasis and drug resistance^17^. Integration of CTC datasets presented us with the opportunity to probe into its validity. For each CTC, we computed two scores indicating the strength of epithelial and mesenchymal phenotypes respectively **(Methods)**. In this analysis, we used tens of canonical markers of each of the concerned phenotypes. We detected near-perfect anti-correlation of (*ρ* = −0.93) the phenotypes across CTCs, coming from all cancer types **(Fig 2-a, Supplementary Fig-3)**. Our findings were consistent when we tracked the association between these phenotypes for CTCs from individual studies **(Supplementary Fig-4)**. Notably, CTC transcriptomes were frequently found on a continuum of epithelial-mesenchymal transition in most of the datasets **(Fig 2-b)**. In selected studies, in spite of being on a continuum, CTCs were found to form clusters towards the epithelial and the mesenchymal poles respectively **(Supplementary Fig-4)**. Melanocytes derive from a highly invasive, multi-potent embryonic cell population called the neural crest. It is suggested that the high degree of plasticity and the aggressiveness of malignant melanoma originate due to the re-activation of the embryonic neural crest program, which is silenced in due course of normal melanocyte differentiation^18^. Unlike the CTCs of most cancer types, circulating melanoma cells were found to be clustered exclusively around the mesenchymal pole of the E-M continuum **(Supplementary Fig-4)**. Our scores correlate well with EMT cell line score proposed by Tan and colleagues^19^ **(Fig 2-c)**. As a secondary validation, we constructed a network incorporating regulations among E and M genes under study **(Methods, Supplementary Fig-5)**. Simulation experiments on this network using Ordinary Differential Equations (ODE) resulted in expression anti-correlation (*ρ* = −0.65) between CDH1 and VIM **(Methods, Fig 2-d, Supplementary Fig-6)**.

**Figure 2.**
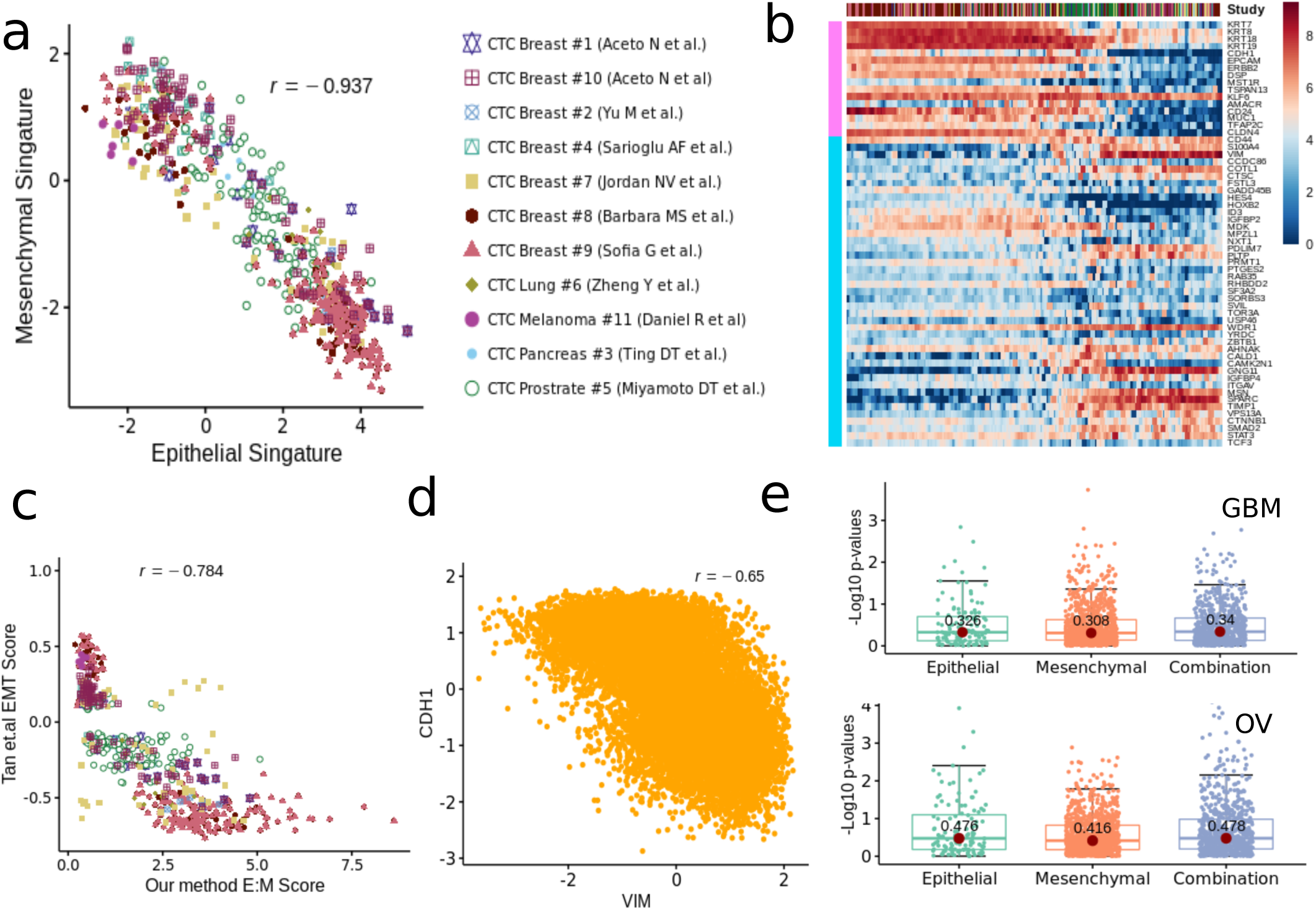
Epithelial-mesenchymal transition in cancer metastasis: (a) Scatter plot showing anti-correlation between epithelial and mesenchymal phenotypes across studies.(b) The moving average smoothen loge(expression+1) of CTC dataset on epithelial and mesenchymal markers where cells are ordered based on the ratio of epithelial and mesenchymal signatures calculated as described in the main methods. (c) Correlation plot of our method E:M score with Tan and colleagues EMT cell line score where negative EMT cell line score corresponds to epithelial like cells and higher E:M score means more epithelial like cells. (d) CDH1 -VIM anti-correlation observed due to simulation of EMT associated regulatory network. (e) Box-plot showing the superiority of the E-M gene pairs, over the E-E and M-M pairs for predicting cancer survival

### Hybrid EMT relates to poor prognosis of the disease

Related to E-M transition, recent *in silico, in vitro*, and *in vivo* studies have indicated that EMT/MET need not be a binary phenomenon, Instead cells may acquire stably one or more hybrid epithelial/mesenchymal (E/M) phenotype(s)^17^. More importantly, these hybrid E/M phenotypes may be more aggressive than cells on either end of the spectrum, due to their enhanced plasticity, increased tumor-initiation potential, resistance to various therapies and anoikis, drug resistance traits, and the ability to migrate collectively to form clusters of CTCs - the key drivers of metastasis^20^. Most of the analysis of EMT has been largely at a bulk level with limited markers^21^, However, individual CTCs can co-express various E and M markers to varying extents, and an increased frequency of hybrid E/M cells correlates with aggressiveness^22, 23^. We performed survival analysis by pairing genes within the E and M sets. To mimic the hybrid phenotype, we also constructed all possible gene pairs across E and M sets **(Methods)**. Across four cancer types (glioblastoma, ovarian, lung, and kidney), E/M gene pairs were found to have higher potential to predict cancer survival relative to the exclusive E or M gene pairs **(Methods, Fig 2-e, Supplementary Fig-7)**.

### Clear patterns observed in expression gradient of immune check-point inhibitor and stemness marker

Loss of major histocompatibility complex (MHC) proteins (aka HLAs) and activation of PD-L1 prevent cytotoxic T cells from attacking tumor cells. Of late, immune checkpoint inhibitors, targeting the PD-1/PD-L1 pathway, have emerged as successful cancer treatment options^24^. In our curated datasets, we found only a minor fraction of CTCs expressing PD-L1. However, PD-L1-MHC anti correlation was evident across studies **(Fig 3-a)**. Two datasets containing the maximum number of PD-L1-activated breast CTCs showed concurrence of PD-L1 with mesenchymal phenotype **(Supplementary Fig-8)**. To date, multiple studies have linked EMT to the formation of cancer stem cells (CSCs). In a seminal paper, Mani and colleagues demonstrated the generation of a CD44^high^/CD24^low^, mammary stem cell-like population due to the induction of EMT. These cells were able to initiate tumors quite efficiently in the mouse. We tracked expression changes in CSC markers along E-M continuum^25^. CD44^high^/CD24^low^ CTCs indeed emerge late in the spectrum, following EMT induction.**(Fig 3-b)** This demonstrates how integrative analysis of CTC transcriptomes can help pinpoint stem-like phenotypes, with high tumorogenesis potential.

**Figure 3.**
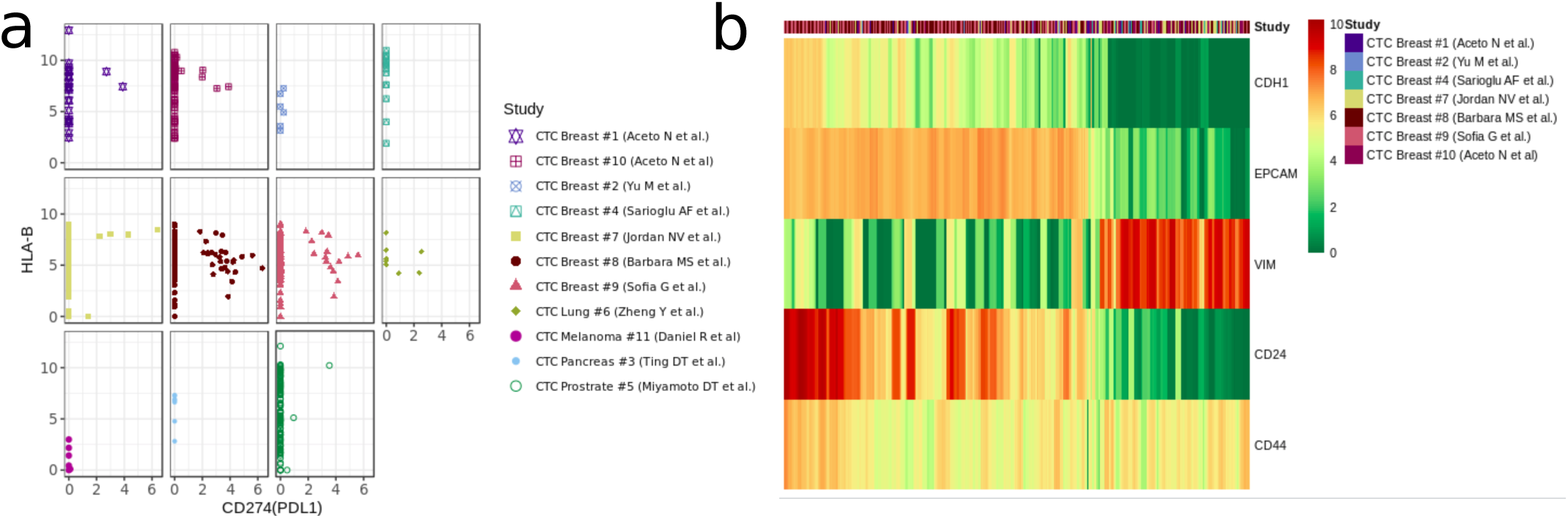
Patterns observed in expression gradient of immune check-point inhibitor and stemness markers. (a) The scatter plot of PDL1 and HLA-B expression in each study. (b) The moving average smoothen loge(expression+1) of specific epithelial, mesenchymal and cancer stem cell markers, across breast CTCs,ordered based on the ratio of epithelial and mesenchymal signatures calculated as described in the main methods.

### CTC-PBMC classification system

We trained a classifier on publicly available single cell expression profiles of human CTCs and PBMCs. Expression datasets curated from independent studies were subjected to rigorous data preprocessing steps **(Methods)**. Notably, the state of the art batch effect removal methods (mnnCorrect^26^ and Seurat^27^) failed to improve the performance of the classification algorithms, compared to a simple median normalisation baseline. We compared the performance of three classifiers - Naïve Bayes^28^, Random Forest^29^, and Gradient Boosting Machine^30^. We evaluated the classifiers by holding out individual CTC and PBMC datasets as test data. We also evaluated the classifiers on all CTC-PBMC data pairs **(Methods)**. Best performance was observed with Naïve Bayes, with a median accuracy recorded at ∼99% and ∼98% respectively **(Fig 4-b,c)**.

**Figure 4.**
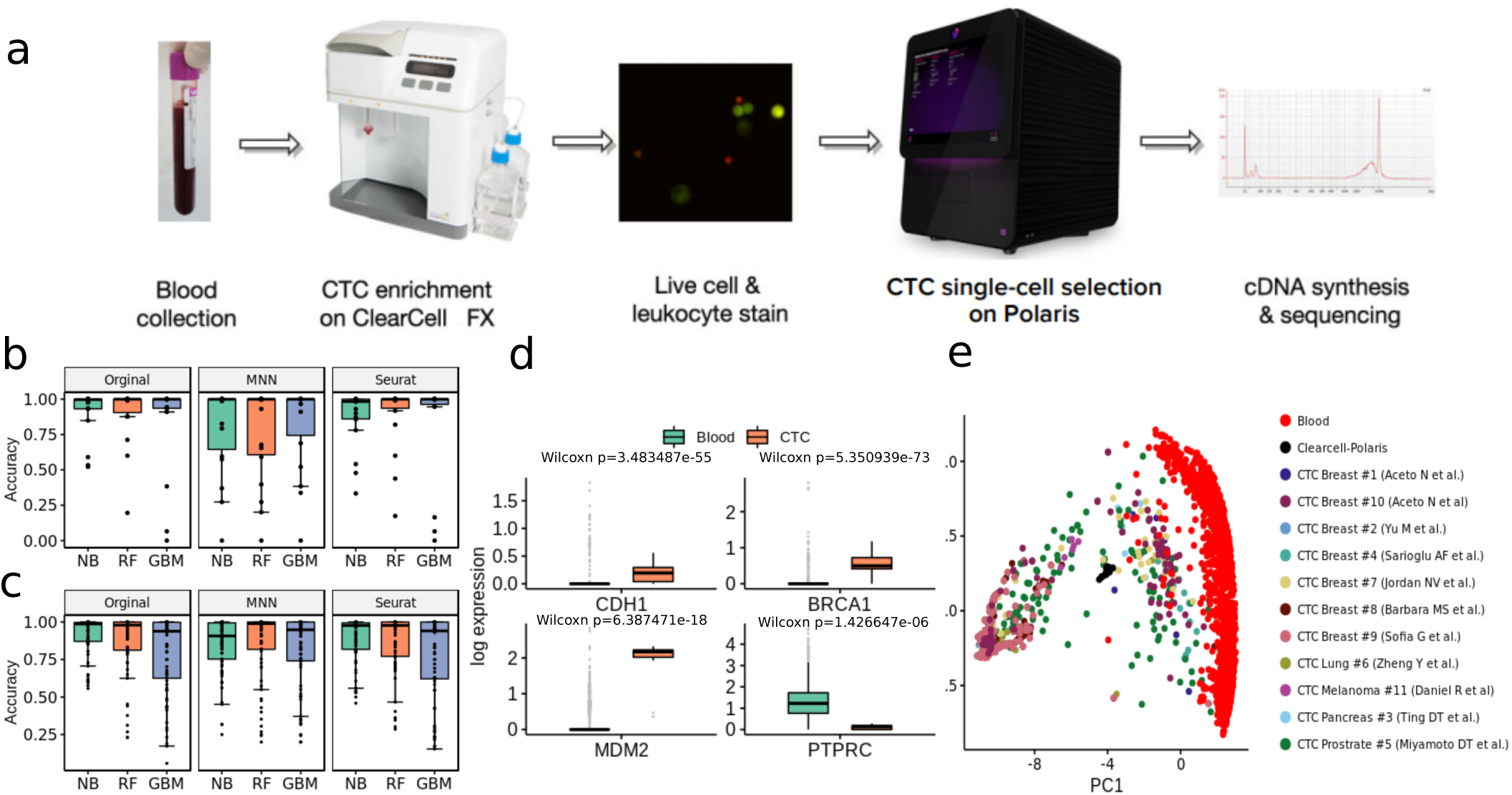
Label free detection and characterisation of CTCs. (a) ClearCell-Polaris workflow involving size-based CTC enrichment by ClearCell FX system, followed by single cell selection and CD45/CD31 depletion using Polaris. (b) Performance of various machine learning algorithms in distinguishing between CTCs and PBMCs. Cells in each dataset were tested against a classifier trained on the remaining datasets. Box plots show the prediction accuracy’s for different choices of classification algorithms (Naive Bayes or NB, Random Forest or RF, Gradient Boosting Machine or GBM) and normalisation/batch-effect correction methods. (c) Box plots showing accuracy’s on held out dataset pairs consisting of a blood and a CTC study.(d) Box-plots showing canonical epithelial/breast cancer specific markers, up-regulated in the CTC population compared to the PBMCs. As expected, PTPRC, a pan leukocyte maker shows elevated expression levels in PBMCs as compared to CTCs. (e) Reference Component Analysis (RCA) based 2D projection of CTCs. PBMCs (red) are visibly separated from CTCs. CTCs enriched using the ClearCell-Polaris workflow cluster with CTCs of other types

### Identification of CTCs captured using novel label-free microfluidic workflow

Existing technologies enrich CTCs with some level of contaminating white blood cells (WBCs). This poses a significant challenge in differentiating CTCs from immune cells. We addressed this challenge by integrating two commercially available microfluidic systems namely Biolidics ClearCell FX System^31^ and the Fluidigm Polaris™ system^16^ **(Methods, Fig 4-a)**. In the proposed workflow CTCs are enriched in two steps - size-based enrichment by ClearCell, followed by CD45 (leukocyte marker) and CD31 (endothelial cell marker) based negative selection by Polaris^16^.

To validate the workflow and the accompanying PBMC-CTC classification system, we processed peripheral blood samples of three HER2-, stage IV breast cancer patients (identified as P3, P4, P5) through the microfluidic device ensemble **(Methods, Supplementary Fig-9)**. Polaris could retrieve 13, 12 and 32 cells from the blood samples of patients P3, P4, P5 respectively. 15 of these 57 cells passed the filtering criteria **(Supplementary Fig-10)**. All 15 cells were classified as CTCs. We used additional validation criteria to determine the carcinogenic origin of the captured cells. When compared to a set of randomly selected PBMCs, ClearCell Polaris captured cells showed elevated expression of breast cancer-specific markers BRCA1 and MDM2^32^ **(Fig 4-d)**. We also detected up-regulation of CDH1, a canonical epithelial cell marker. Expression of CD45 (PTPRC) was considerably low in these cells compared to the PBMC transcriptomes **(Fig 4-d)**. Reference component analysis (RCA) allows noise-free single cell clustering, by projecting single cell transcriptomes on reference bulk expression data. We subjected all CTC and PBMC transcriptomes to RCA analysis^33^. ClearCell-Polaris captured CTCs grouped with other CTCs, whereas the PBMCs formed a separate cluster **(Methods, Fig 4-e, Supplementary Fig-11)**.

## Discussion

CTCs have been shown to be of prognostic significance in patients with various cancers^2, 15, 34^. It is suspected that a large number of CTCs do not portray the signature of cancer epithelium, largely due to their acquired phenotype that is suitable for migration^34^. The proposed machine learning based bioinformatics approach accurately distinguishes CTCs from regular immune cell sub-types. This is achieved by the integration of publicly available CTC datasets and machine learning based model training. We provide a user-friendly R package for CTC classification that provides a probabilistic score indicating potential carcinogenic origin of individual cells. Our reported ClearCell^®^ Polaris™ workflow, in tandem with the machine learning based CTC-immune cell classification system, for the first time, enables truly unbiased detection of circulating tumor cells. With declining per cell cost associated with single-cell gene expression screening, we speculate a high adoption rate for our proposed approach.

Integrative study of CTC transcriptomes presented us with the opportunity to discover consistent pan-cancer CTC surface-proteins, besides EpCAM. We looked for surface-protein coding genes which are deferentially upregulated in CTCs over blood cells **(Supplementary Note-5)**. Most remarkable among these were ERBB3, LTBP1, TACSTD2 and EFNA1 **(Supplementary Fig-12)**. In addition to EpCAM, some of these markers might be useful to broad-base marker dependent capture of CTCs.

## Methods

### Description of datasets

We collected single-cell RNA-seq (scRNA seq) data of circulating tumor cells (CTCs) and peripheral blood mononuclear cells (PBMCs) from 15 different studies^2, 10, 15, 34–40, 40–42^ **(Supplementary Table-1)**. We acquired 729 single CTCs from 11 of these 15 studies. On the other hand, 6 of these studies supplied a total of 37107 PBMCs. Two studies provided both CTCs and PBMCs. The CTC data entailed five cancer types breast, prostate, melanoma, lung, and pancreas. Notably, circulating breast tumor cells in the data were supplied by seven different studies. Remaining cancer types were represented by single studies.

### Data Pre-processing

We downloaded raw read count data for every study from their respective sources **(Supplementary Table 1)**. We also considered 15 CTC transcriptomes with exonic read count >50000, from three HER2-breast cancer patients (details given below). While merging, we found 15043 genes common across all the datasets. First, we discarded the poor quality cells that had less than 6% of the genes having non zero expression. The filtering step retained about 35% (13235) of the input cells. Genes with read count ≥5 in at least 10 cells were retained. A total of 12624 genes were left after this. Among the 13235 cells, 737 were CTCs. In the remaining 12498 PBMCs, one single study (EGAS00001002560)^40^ alone supplied 11697 cells leading to the class-imbalance problem. We decided to retain cells having total read counts ≥5000. Our final data contained a 12624 expressed genes and 3079 cells, of which 729 were CTCs. At this stage, we standardized the library depths using median normalization. The expression matrix thus obtained was *log*_*e*_ transformed after addition of 1 as pseudo-count. Different gene selection techniques used for the various downstream analyses are mentioned in the subsequent sections.

### Construction of epithelial and mesenchymal signatures

While integrating, we found 17609 genes common across 729 CTCs coming from 11 publicly available CTC datasets. After applying the cell and gene filtering steps (as discussed above), we were left with an expression matrix consisting of 14027 genes and 722 CTCs. We constructed a panel of 180 well-known epithelial, mesenchymal, and cancer stem cell markers combining information from the CellMarker database^43^ and existing literature. The expression matrix of marker genes thus obtained was subjected to stricter criteria for gene and cell selection. We retained 718 cells that expressed at least 10% of these marker genes. Marker genes having minimum read count >5 in at least 30% of these cells were selected for the subsequent analyses. The resulted matrix consisted of 718 cells and 86 marker genes (16 epithelial, 43 mesenchymal, and 27 cancer stem cell markers, see **(Supplementary Table 2**). We normalized and log-transformed the matrix as mentioned above. For each cell, we computed a comprehensive score for both epithelial and mesenchymal phenotype. To compute the score we first applied Z-score transformation on each cell. To create the signature for specific phenotype, for each cell we combined Z-transformed marker expressions using the below formula.

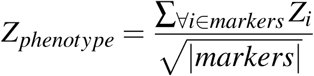

Here *Z*_*phenotype*_ is a comprehensive phenotype specific score computed over individual Z-transformed marker expressions denoted by *Z*_*i*_, where *markers* denotes the set of markers corresponding to the concerned phenotype.

### Simulation of E-M continuum

We identified the regulatory interactions among epithelial (E) and mesenchymal (M) genes under study, together with their connections to canonical regulators of EMT and MET such as the double negative feedback loops involving miR-200, ZEB and GRHL2 **(Supplementary Note-3)**. For the constructed network, an ensemble of mathematical models was then created using RACIPE (RAndom CIrcuit PEr-turbation), which considers a set of kinetic parameters randomly chosen from within the biologically relevant ranges^44^. This helps to identify the robust gene expression signatures that can emerge due to a given network topology. The simulations were performed in triplets to avoid numerical artifacts/variations due to random sampling. Such ensemble of models are usually based on ordinary differential equations (ODEs), such as the one mentioned below.

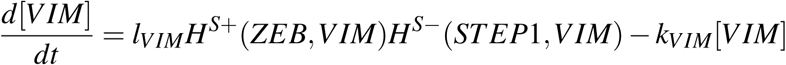

where [*VIM*] is the concentration of VIM, and *l*_*VIM*_ and *k*_*VIM*_ are its production and degradation rates respectively. *H*^*S*+^(*X,Y*)/ *H*^*S*−^(*X,Y*) are the shifted Hill functions that result in up-regulation/down-regulation caused in the expression of Y due to X.

### Survival analysis for hybrid E/M phenotype

We investigated the clinical relevance of the E, M, and hybrid E/M phenotypes by leveraging those in patient survival prediction for four cancer types from The Cancer Genome Atlas (TCGA) project^45^. We focused our analyses on four TCGA cancer types with high-quality overall survival data i.e kidney renal clear cell carcinoma (KIRC), glioblastoma multiforme (GBM), ovarian serous cyst adenocarcinoma (OV) and lung squamous cell carcinoma (LUSC)^46^. We only used data of the patients for whom the survival information was available. Raw read count data of TCGA samples were extracted from the Recount2^47^ repository. The dataset corresponding to each cancer type was median-normalized and *log*_*e*_ transformed after addition of 1 as pseudo count. We used all 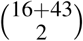 possible pairs of 16 epithelial and 43 mesenchymal markers, one at a time to divide the patient samples into two groups. For each gene in a pair, Z-scores were computed across the patient samples. Stouffer’s Z was computed for each gene pair by combining the Z-scores, computed independently. For every cancer, patient groups were formed by applying a cutoff at the median of the Stouffer’s Z score vector. Survival curves thus obtained were compared using the log-rank test. We used survminer R package^48^ for the survival analyses **(Supplementary Table 3)**.

### Classification of cancer and blood transcriptomes

To model the phenotypic identities of CTCs and PBMCs, we trained various classification models. To broad-base our feature selection we used about 3000 cell-type specific markers **(Supplementary Table-4)** reported in the CellMarker database^43^. Besides, median normalization, we subjected the data to two different batch correction methods - mnnCorrect^26^ and canonical correlation based batch correction method from the Seurat R package^27^ **(Supplementary Note-1,2)**. We used the h2o APIs^49^ of three popular classification techniques - Naive Bayes (NB)^28^, Gradient Boosting Machines (GBM)^30^ and Random Forest (RF)^29^. To evaluate the model generalisability we performed two different experiments. In the first experiment, we held out each dataset to assess the model trained on the remaining datasets. In the second experiment, we tested the model by holding out all the possible combinations of one CTC and one PBMC data. Besides the accuracy percentage, we reported additional model evaluation metrics such as F1 score, Mathews correlation coefficient (MCC) and Cohen’s kappa as applicable **(Supplementary Table-5,6)**.

### Sample collection

Blood specimens of three HER2-breast cancer patients (identified as P3, P4, P5) were obtained from the National Cancer Center Singapore, with informed consent in accordance with the approved procedures under the institutional review board (IRB) guidelines (CIRB no. 2014/119/B). The clinical sample collection protocols were reviewed and approved by the Sing Health Centralised Institutional Review Board. All three subjects had ER+/PR+/HER2-hormone receptor status as analyzed by immunohistochemistry. The determination of estrogen receptor (ER), progesterone receptor (PR) and human epidermal growth factor receptor 2 (HER2) status by immunohistochemistry in this study was based on the latest recommendations of the American Society of Clinical Oncology and the College of American Pathologists. For P3, blood was drawn (baseline) in August, 2016 for CTC enrichment. Following this P3 was on chemotherapy. P4 and P5 were on chemotherapy before their blood samples were collected for CTC enrichment in August and September of 2016, respectively.

### CTC enrichment

Blood samples were collected in 9 mL K3EDTA blood collection tubes (Greiner Bio-One, 455036). 6 – 8.5 mL of whole blood was processed for each run. Red blood cells were first removed with the addition of red blood cell (RBC) lysis buffer (G-Bioscience, St. Louis, MO, USA) and incubation for 10 minutes at room temperature. Lysed RBCs in the supernatant were discarded after centrifugation. The nucleated cell pellet was suspended in a ClearCell re suspension buffer prior to CTC enrichment on ClearCell FX system (Biolidics Limited)^31^, performed in accordance with manufacturer’s instructions.

### Immunofluorescence suspension staining

The enriched CTC blood sample was centrifuged at 300 g for 10 min and concentrated to 70 μL. The cells were stained with the addition of the following markers and antibodies for 1 hour: CellTracker Orange (CTO) (Thermo Fisher, C34551), Calcein AM (Thermo Fisher, L3224), CD45 antibody-conjugated with Alexa 647 (Bio Legend, 304020), and CD31-conjugated with Alexa 647 (Bio Legend, 303111). 15 μL of RPMI with 10% FBS (Gibco) and 3 μL of RNase inhibitor (Thermo Fisher, N8080119) were also added to improve the viability and RNA quality of the cells. After incubation, 13 mL of PBS was added to dilute the staining reagents. The sample was spun down at 300 g for 10 min and concentrated to 45 μL. In order to achieve optimal buoyancy in an integrated fluidic circuit (IFC), 45 μL of CTCs was mixed with 30 μL Cell suspension Reagent (Fluidigm, 101-0434) to achieve 75 μL of cell mix.

### Integrated Fluidic Circuit (IFC) operation

The Polaris IFC is first primed using Polaris system (Fluidigm)^16^ to fill the control lines on the fluidic circuit, load cell capture beads, and block the inside of PDMS channels to prevent non-specific absorption/adsorption of proteins. To capture and maintain the single cells in the sites, the capture sites (48 sites) are preloaded with beads that are linked on-IFC to fabricate a tightly packed bead column during the IFC prime step. After completion of the prime step, the cell mix (cells with suspension reagent) is loaded in three inlets (25 μL each of cell mix) on the Polaris IFC and single cells with CTO+ Calcein AM+ CD45-CD31-are selected to capture sites. Finally, the single cells are processed through template-switching mRNA-seq chemistry for full-length cDNA generation and preamplification on-IFC.

### mRNA-seq library preparation and sequencing

SMARTer® Ultra® Low RNA Kit for Illumina® Sequencing (Clontech®, 634936) was used to generate preamplified cDNA. The selected and sequestered single cells were lysed using Polaris cell lysis mixture. The 28-μL cell lysis mix consists of 8.0 μL of Polaris Lysis Reagent (Fluidigm, 101-1637), 9.6 μL of Polaris Lysis Plus Reagent (Fluidigm, 101-1635), 9.0 μL of 3 SMART™ CDS Primer II A (12 M, Clontech, 634936), and 1.4 μL of Loading Reagent (20X, Fluidigm, 101-1004). The thermal profile for single-cell lysis is 37°C for 5 min, 72°C for 3 min, 25°C for 1 min, and hold at 4°C. The 48-μL preparation volume for reverse transcription (RT) contains 1X SMARTer Kit 5X First-Strand Buffer (5X; Clontech, 634936), 2.5-mM SMARTer Kit Dithiothreitol (100 mM; Clontech, 634936), 1-mM SMARTer Kit dNTP Mix (10 mM each; Clontech, 634936), 1.2-μM SMARTer Kit SMARTer II A Oligonucleotide (12 μM; Clontech, 634936), 1-U/μL SMARTer Kit RNase Inhibitor (40 U/μL; Clontech, 634936), 10-U/μL SMARTScribe™ Reverse Transcriptase (100 U/μL; Clontech, 634936), and 3.2 μL of Polaris RT Plus Reagent (Fluidigm, 101-1366). All the concentrations correspond to those found in the RT chambers inside the Polaris IFC. The thermal protocol for RT is 42°C for 90 min (RT), 70°C for 10 min (enzyme inactivation), and a final hold at 4°C.

The 90-μL preparation volume for PCR contains 1X Advantage 2 PCR Buffer [not short amplicon (SA)](10X, Clontech, 639206, Advantage® 2 PCR Kit), 0.4-mM dNTP Mix (50X/10 mM, Clontech, 639206), 0.48-μM IS PCR Primer (12 μM, Clontech, 639206), 2X Advantage 2 Polymerase Mix (50X, Clontech, 639206), and 1X Loading Reagent (20X, Fluidigm, 101-1004). All the concentrations correspond to those found in the PCR chambers inside the Polaris IFC. The thermal protocol for preamplification consists of 95°C for 1 min (enzyme activation), five cycles (95°C for 20 s, 58°C for 4 min, and 68°C for 6 min), nine cycles (95°C for 20 s, 64°C for 30 s, and 68°C for 6 min), seven cycles (95°C for 30 s, 64°C for 30s, and 68°C for 7 min), and final extension at 72°C for 10 min. The preamplified cDNAs are harvested into 48 separate outlets on the Polaris IFC carrier. The cDNA reaction products were then converted into mRNA-seq libraries using the Nextera® XT DNA Sample Preparation Kit (Illumina, FC-131-1096 and FC-131-2001, FC-131-2002, FC-131-2003, and FC-131-2004) following the manufacturer’s instructions with minor modifications. Specifically, reactions were run at one-quarter of the recommended volume, the tagmentation step was extended to 10 min, and the extension time during the PCR step was increased from 30 to 60 s. After the PCR step, samples were pooled, cleaned twice with 0.9× Agencourt AMPure XP SPRI beads (Beckman Coulter), eluted in Tris + EDTA buffer and quantified using a high-sensitivity DNA chip (Agilent). The pooled library was sequenced on Illumina MiSeq™ using reagent kit v3 (2×75 bp paired-end read). The sequencing data generated was processed by standard bioinformatic pipeline **(Supplementary Note 4)**.

### Reference component analysis of CTCs and PBMCs

For reference component analysis (RCA), we used the global panels supplied as part of the RCA R package^33^. Each of the global panels consisted of numerous tissue samples. RCA^33^ uses cell type specific genes for measuring correlation between the tissue types and the input single cells. Due to low amount of starting RNA, single cell expression data is far noisier than bulk expression data. As a result, tissue types represented by lowly expressed feature genes can potentially give rise to significant levels of noise. In each global panel, we therefore retained 50% of the tissue types with highest median expression of the feature genes. RCA^33^ analysis provided us with both single cell - tissue correlation heat-map and 2D projection of the individual transcriptomes.

### Data and Code Availability

The data-set used in the study are available from links mentioned in the Supplementary Table-1. Single cell sequencing data generated for this paper is deposited at GEO with accession number GSE129474 [Token: qdkvyayyprwjvix]. Code used for analysis is available at this link and a R package is available at link.

## Acknowledgements

This work is partially supported by the INSPIRE Faculty Grant (DST/INSPIRE/04/2015/003068) awarded to D.S. by the Department of Science and Technology (DST), Govt. of India. M.K.J is supported by Ramanujan Fellowship provided by SERB, DST, Government of India (SB/S2/RJN-049/2018).

## Author contributions statement

DS and NR conceived the project. AI and KG performed the majority of the analyses under the supervision of DS. SS assisted AI and KG in the computational analyses. MKJ planned the EMT modeling. KH, BS, VS performed the associated analysis under MKJ’s supervision. NR, JW, AAB conceived integration of FX and Polaris. NR and YFL developed the label-free workflow. YSY provided the patient samples. YFL tested patient samples and NeR assisted NR in data analysis. All the authors discussed the results, co-wrote and reviewed the manuscript.

## Additional information

