## Supplementary Information for "Integrative analysis and machine learning based characterization of single circulating tumor cells"

### Supplementary Note 1: Integration of dataset using the mnnCorrect(scran) method.

For performing the mutual nearest neighbors method we processed the median normalized CTC+Blood dataset to $\log_e$ transformation after addition of 1 as a pseudo count. mnnCorrect()[1] function from scran package with k=5 was used to perform the correction of the integrated CTC+Blood dataset. The corrected data returned by the function was used for further analysis. For machine learning, purpose mnnCorrect[1] was run twice first to correct the train data and then to correct the test data with the corrected train data.

### Supplementary Note 2: Integration of dataset using the CCA (Seurat) method.

For performing the canonical correlation method as described by Butler et.al[2] paper we followed the standard pipeline :

1. Each study raw count data obtained from gene filtering step (Main methods) was converted into a Seurat R object.
2. Each study then was subjected to default Seurat pipeline of normalization, finding variable genes and scaling of the data.
3. After having the Seurat R object for each study they were subject to RunMultiCCA() / RunCCA() function with Number of canonical vectors set to 2.
4. After the CCA the dataset was aligned to canonical vectors by using AlignSubspace() function.

For machine learning, the training data was first aligned as mentioned in the steps above and stored as one Seurat R object. This Seurat R data was again subjected to the RunMultiCCA() / RunCCA() function along with test data to obtain the final and train dataset.

### Supplementary Note 3: Network analysis to investigate the mechanistic basis of EMT continuum phenotype observed in the data analysis.

To investigate the mechanistic basis of our data analysis from multiple CTC datasets, we explored the expansive literature for a functional implication of the genes that were identified in epithelial and mesenchymal signatures for their roles in EMT and/or MET (see Supplementary Table 2). We next constructed a network based on the functional implications of these genes in EMT and/or MET, including their effects on the regulatory feedback loops involving miR-200, ZEB and GRHL2 – the fulcrum of epithelial-mesenchymal plasticity. We mapped the connections known to promote EMT and metastasis (SNAIL, TIMP1, etc.) through promoting ZEB and/or inhibiting miR-200. Moreover, some genes in this list have already been known to have a direct effect on CDH1 and VIM. Please note that many of the molecules identified here are markers of epithelial and mesenchymal state, and their functional impact on EMT or metastasis remains elusive, hence they were excluded from the network. CDH1 and VIM were chosen to denote epithelial and mesenchymal states respectively; their expression levels are considered as the outcome for the network.

As the data were collected across multiple cancer types, the parameters for each connection are very likely to vary. Hence, instead of applying a single parameter set to the network, we sampled the parameter space via a uniform distribution using a tool called RAndom

CIrcuit PErturbation (RACIPE)[3]. As the name suggests, RACIPE[3] chooses the parameter space of a given circuit randomly to elucidate the robust dynamical outcomes of the network and pinpoint the gene expression signatures most likely to emerge from the given network topology.

To understand the significance of the network topology, we generated random network topologies by swapping the edges while maintaining the degree of each node and the number of activating and inhibiting edges in the network, ensuring the conservation of the nature of the nodes. Furthermore, we characterized the effect of single edge perturbations (SEP's), i.e., change in the sign of one edge in the network at a time, on the correlation between ZEB-miR200 and VIM-CDH1 pairs. With randomized networks (Supplementary Fig-7a,Supplementary Fig-7a), we observe that the correlation between the markers VIM-CDH1 as well as the core elements ZEB-miR200 is strongly negative in the original network, denoted by wildtype (WT). A very small fraction of randomly generated topologies have equal or stronger correlation than WT for both (CDH1, VIM) and (ZEB, miR-200), suggesting the importance of the particular network topology for the observed behavior. Similarly, we observe that most SEPs do not show as a strong correlation in terms of (CDH1, VIM) or (ZEB, miR-200) as the original network (WT, shown in red). The effect of perturbations on the network is calculated by applying a distance metric, Jensen-Shannon Divergence (JSD)[4], on the steady-state frequencies of the networks.

### Supplementary Note 4: Gene expression quantification of CTCs detected by the ClearCell Polaris workflow

An index for RNA-Seq by expectation maximization (RSEM) was generated based on the hg19 RefSeq transcriptome downloaded from the UCSC Genome Browser database. Read data were aligned directly to this index using RSEM/bowtie. Quantification of gene expression levels in counts for all genes in all samples was performed using RSEM v1.2.4[5]. Genomic mappings were performed with TopHat 2 v2.0.13[6], and the resulting alignments were used to calculate genomic mapping percentages. Raw sequencing read data were aligned directly to the human rRNA sequences NR_003287.1 (28s), NR_003286.1 (18S) and NR_003285.2 (5.8S) using bowtie 2 v2.2.4[7], and the percentage of reads aligned to rRNA was then calculated as reads aligned to these sequences divided by the total reads.

### Supplementary Note 5: Exploration of novel surface markers for CTCs.

We performed Wilcoxon's rank-sum test for determining differentially expressed genes between CTCs and blood cells. P-values thus obtained were subjected to multiple test corrections using the Benjamin-Hochberg method (p.adjust function in R). We applied an FDR cut off of 0.05 for selecting the differential genes (DE). DE genes that were expressed in at least 70% of the CTCs were retained. We downloaded a list of the surface proteins from the Cell Surface Protein Atlas (CSPA) database[8] and took intersection with the narrowed set of DE genes. Supplementary Fig-10 displays the selected markers in the order of the gene-wise fold change values.

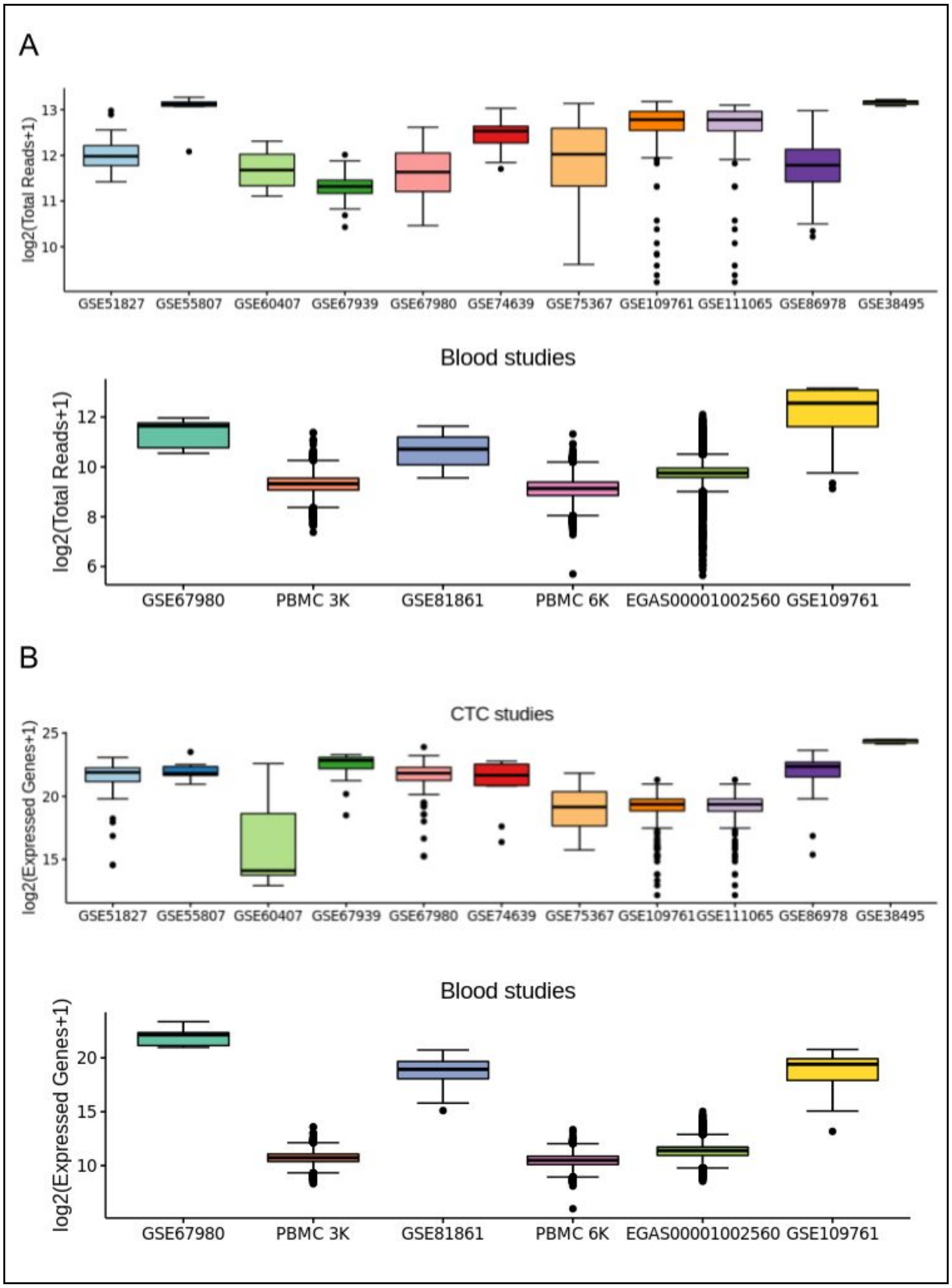


**Supplementary Figure-1:** A) Boxplots show the distribution of total read counts across cells in each dataset. B) Boxplots show the distribution of the number of detected (non zero) genes in each dataset.

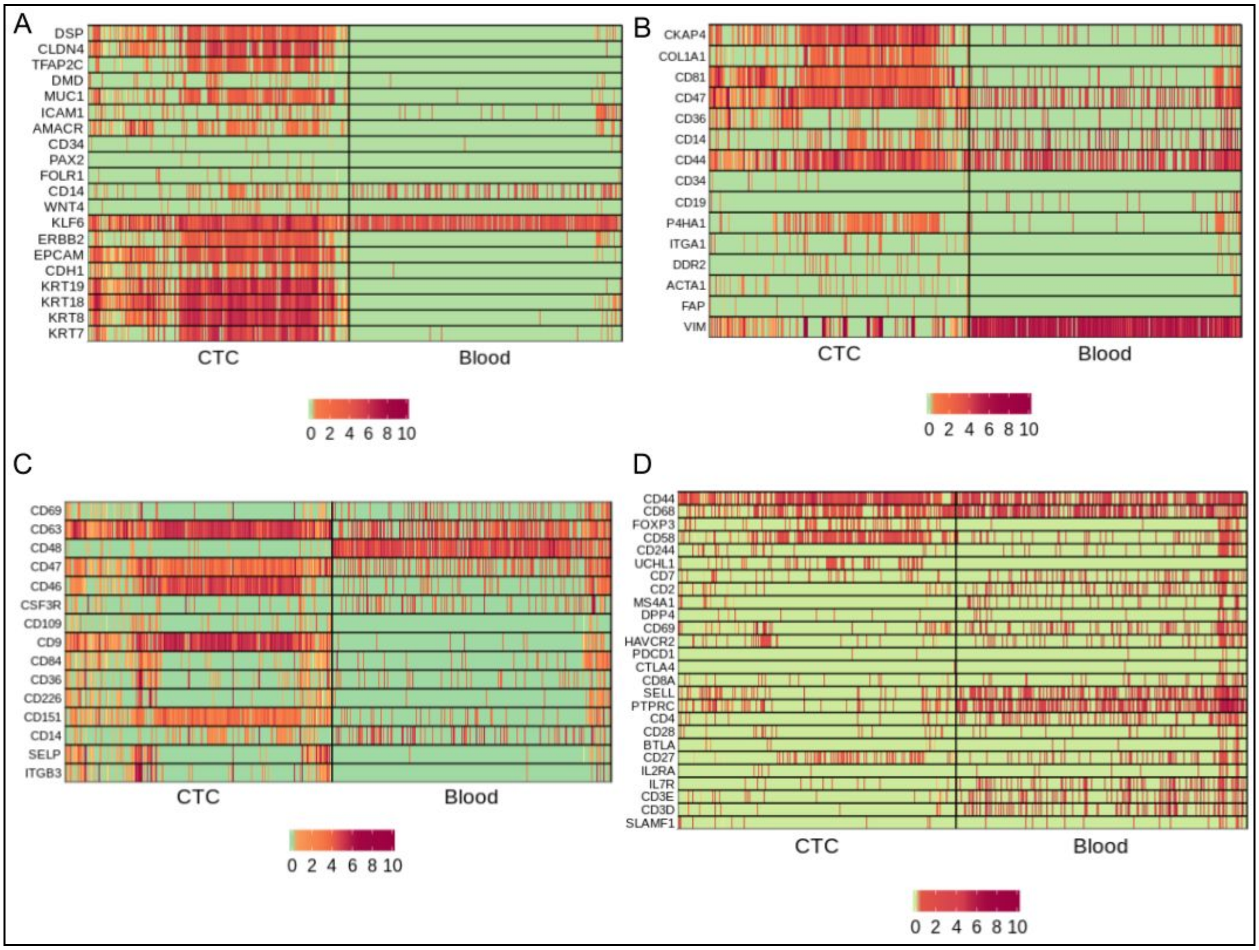


**Supplementary Figure-2:** Expression of known markers in curated CTCs and PBMCs
A) Expression of Epithelial markers in the integrated dataset of CTCs and PBMCs (blood). B) Expression of Fibroblast markers C) Expression of Platelet markers. D) Expression of T-cell markers.

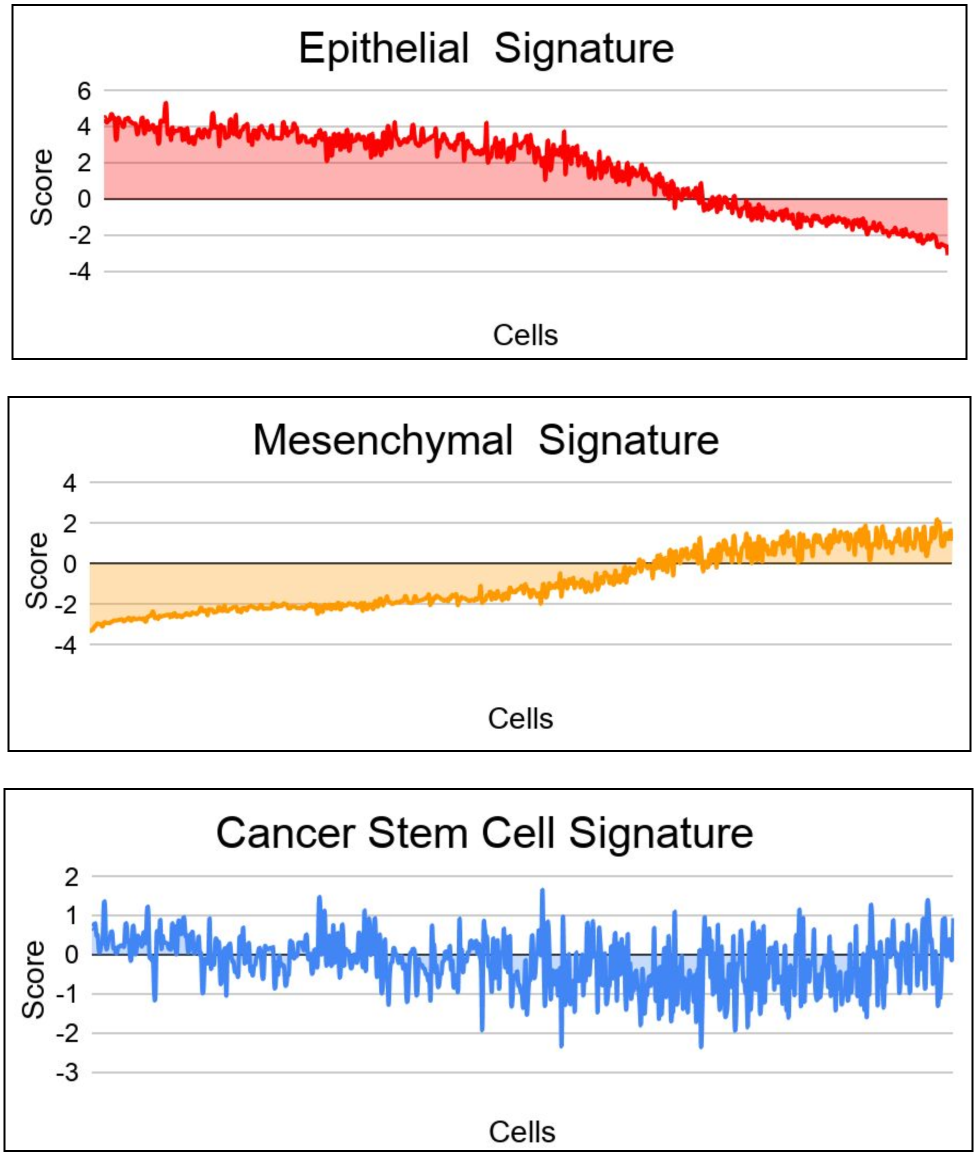


**Supplementary Figure-3:** Combined epithelial, mesenchymal and cancer stem cell signatures.

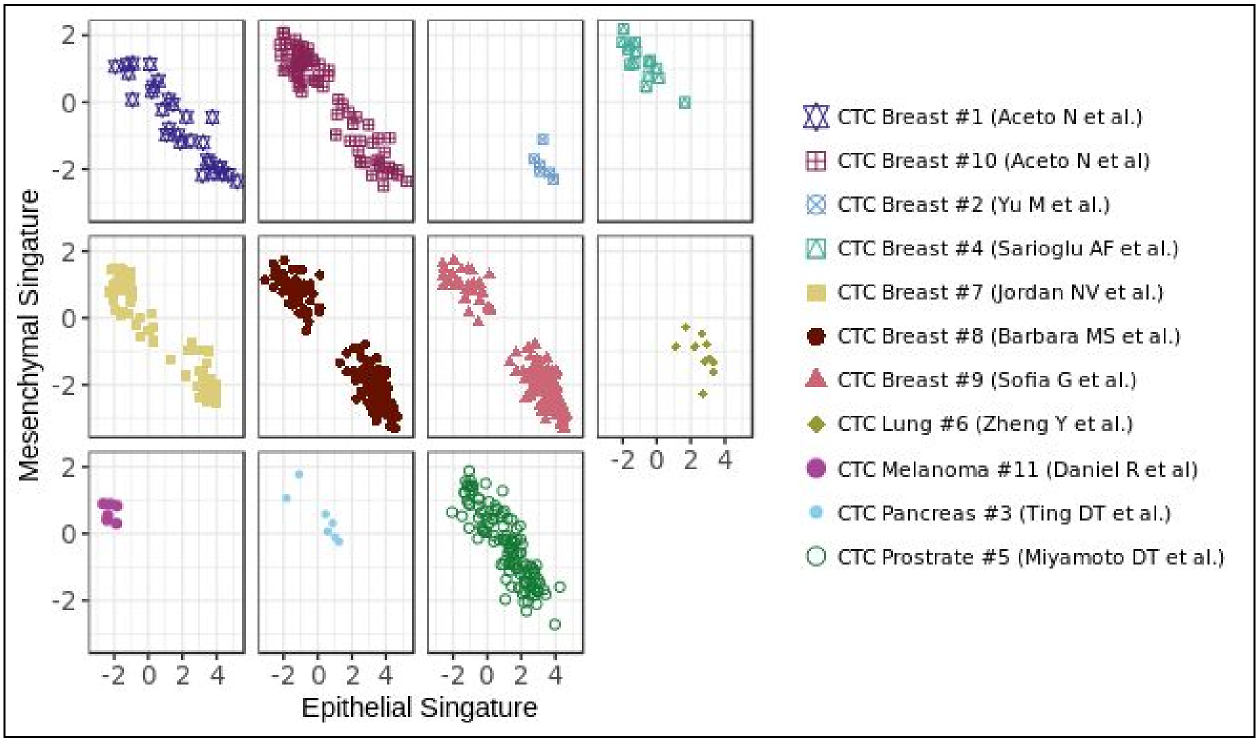


**Supplementary Figure-4:** Scatter plots show Epithelial-Mesenchymal anti-correlation for individual datasets.

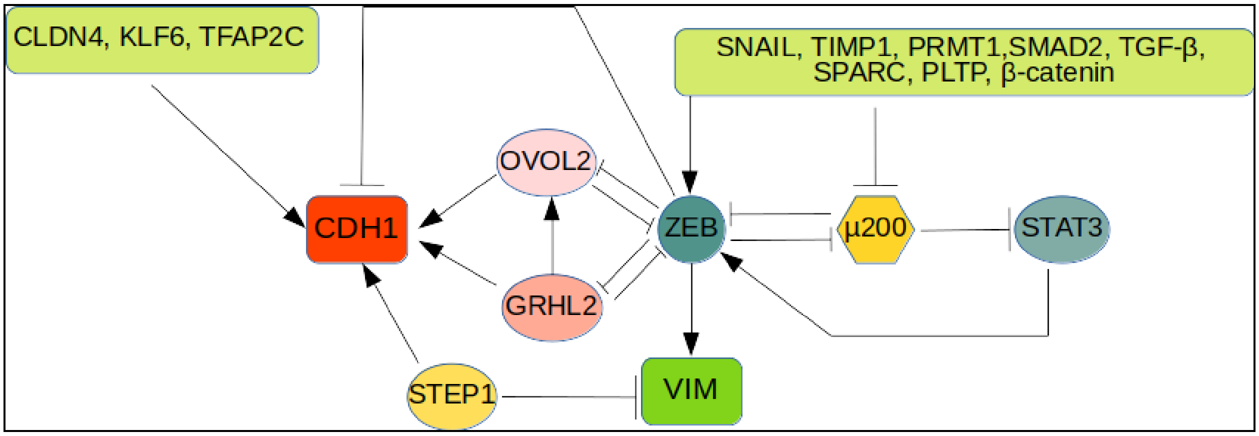


**Supplementary Figure-5:** The network simulated using RACIPE, including genes used for the creation of epithelial and mesenchymal signatures. Here µ200 represents miRNA-200.

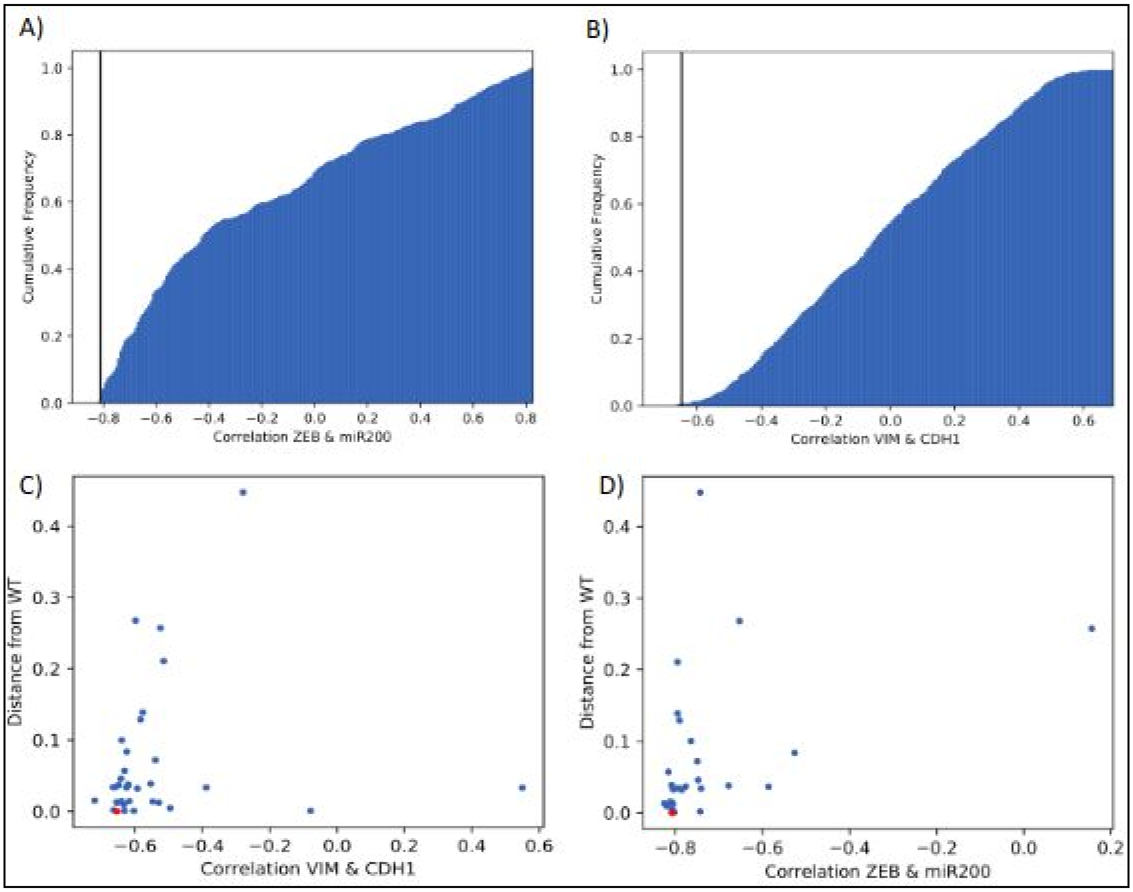


**Supplementary Figure-6:** Top: Cumulative frequency distribution of the correlation between A) VIM and CDH1 and B) ZEB and miR200. The black line represents the correlation coefficient for 'wildtype' (WT) network. Bottom: scatter plots of JSD distance against the correlation between C) VIM and CDH1 and D) ZEB and miR200 for SEP's. Y-axis represents the distance of each SEP from WT (JSD).

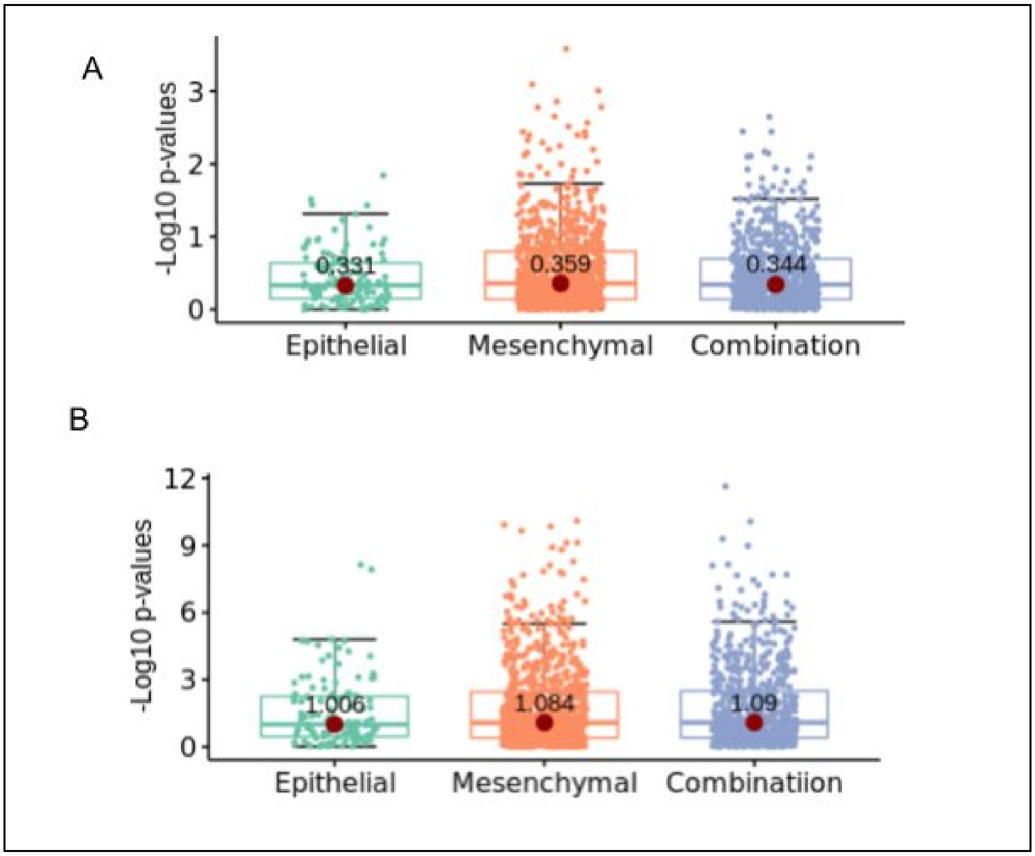


**Supplementary Figure-7:** Hybrid E/M survival analysis on lung (LUSC) and kidney cancer (KIRC) data (see Online Methods for further details)

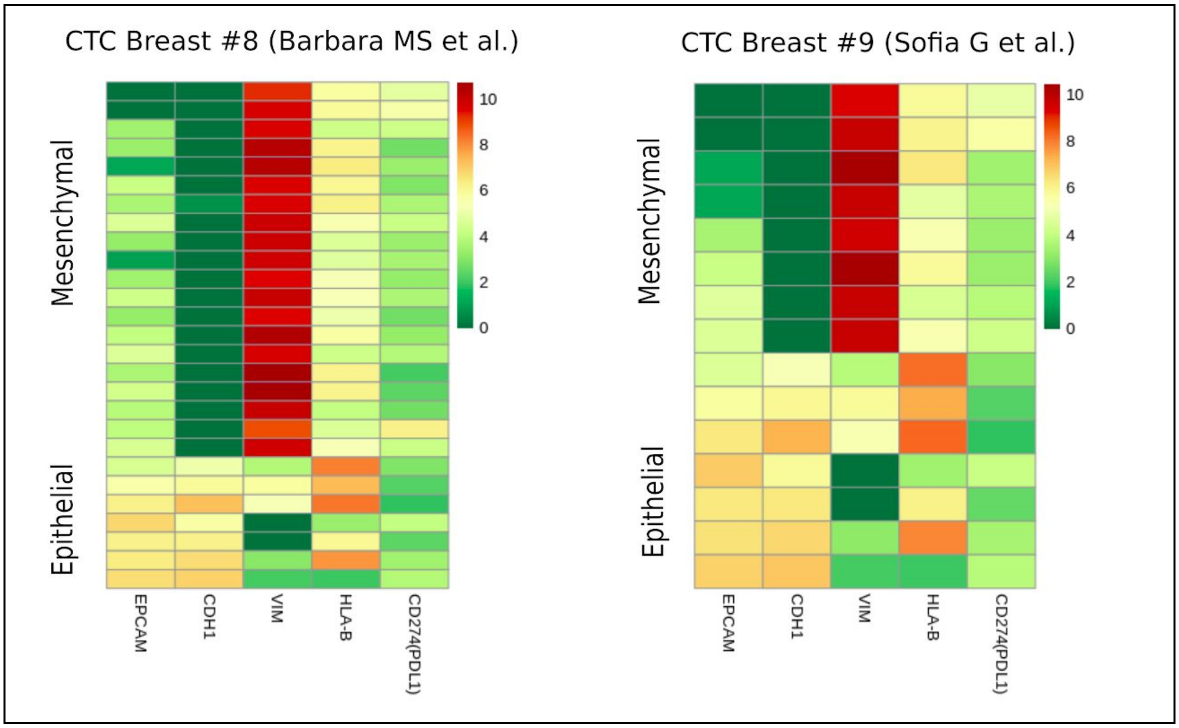


**Supplementary Figure-8:** The heatmap of $\log_e$ (expression+1) of selected epithelial, mesenchymal marker along with PDL1 and HLA-B for cells with non zero PDL1 expression from two specific studies (having maximum numbers of PDL1 expressing cells). Note that we only showed HLA-B expression since this was most well expressed.

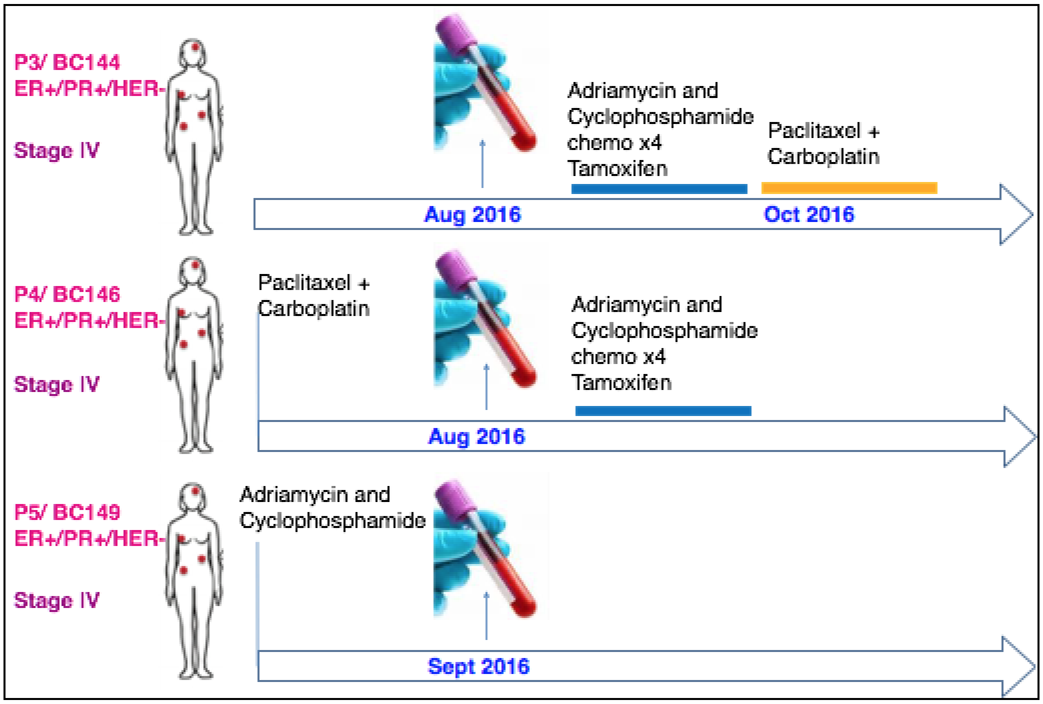


**Supplementary Figure-9:** Treatment history of the patients

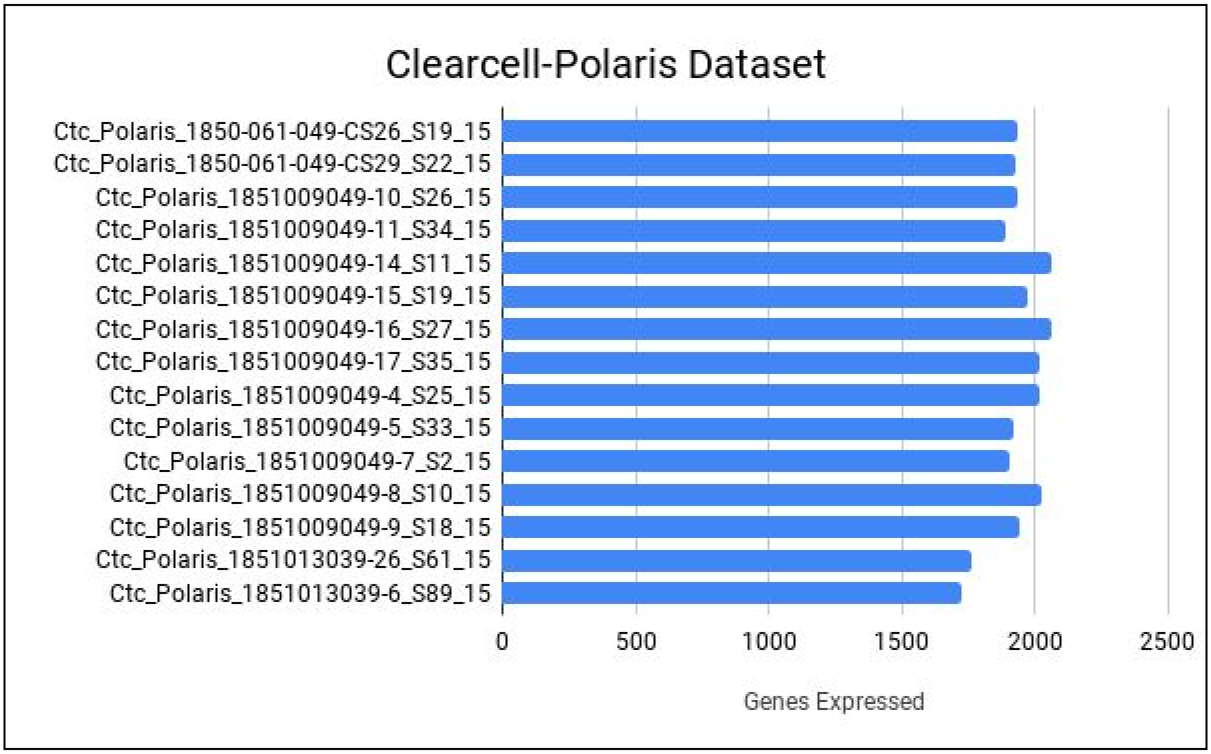


**Supplementary Figure-10:** Number of expressed genes in CTCs detected using the Clearcell-Polaris workflow

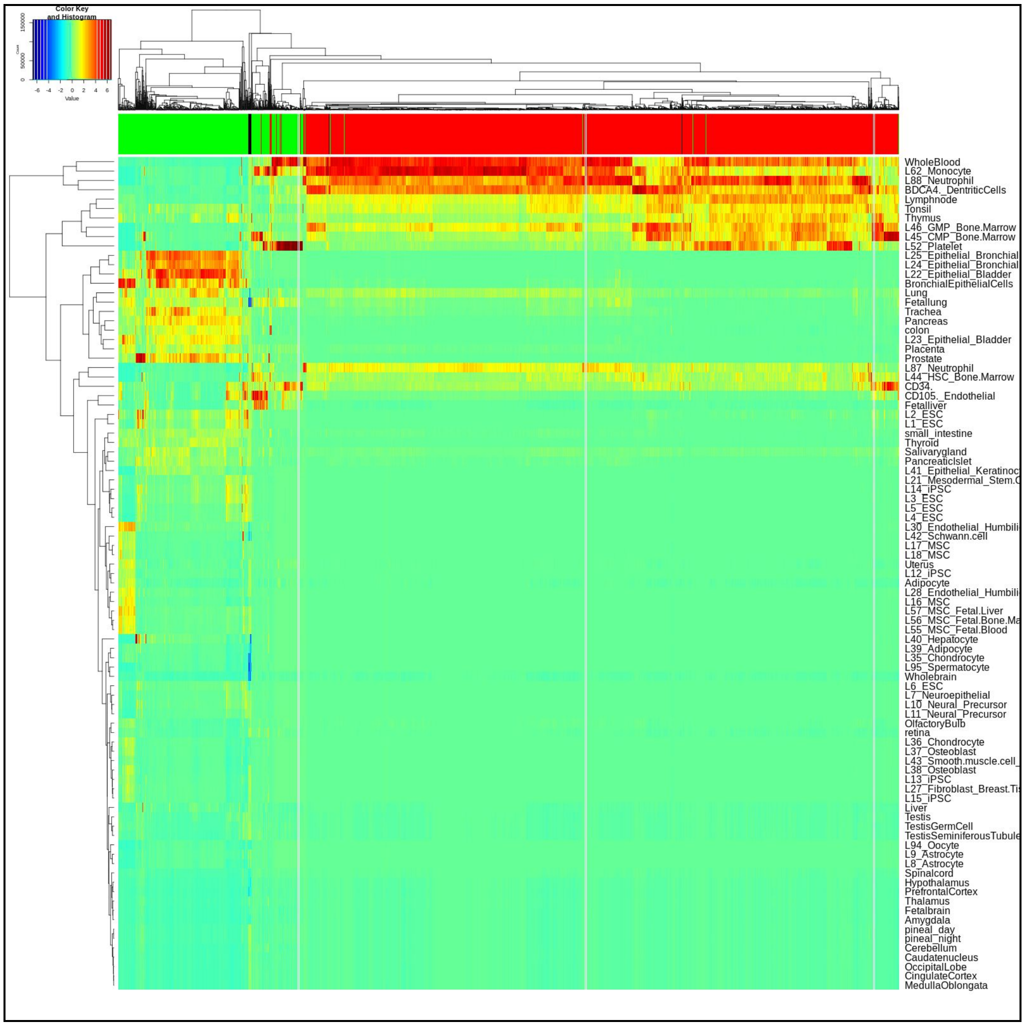


**Supplementary Figure-11**: Tissue - single cell correlation plot obtained from RCA

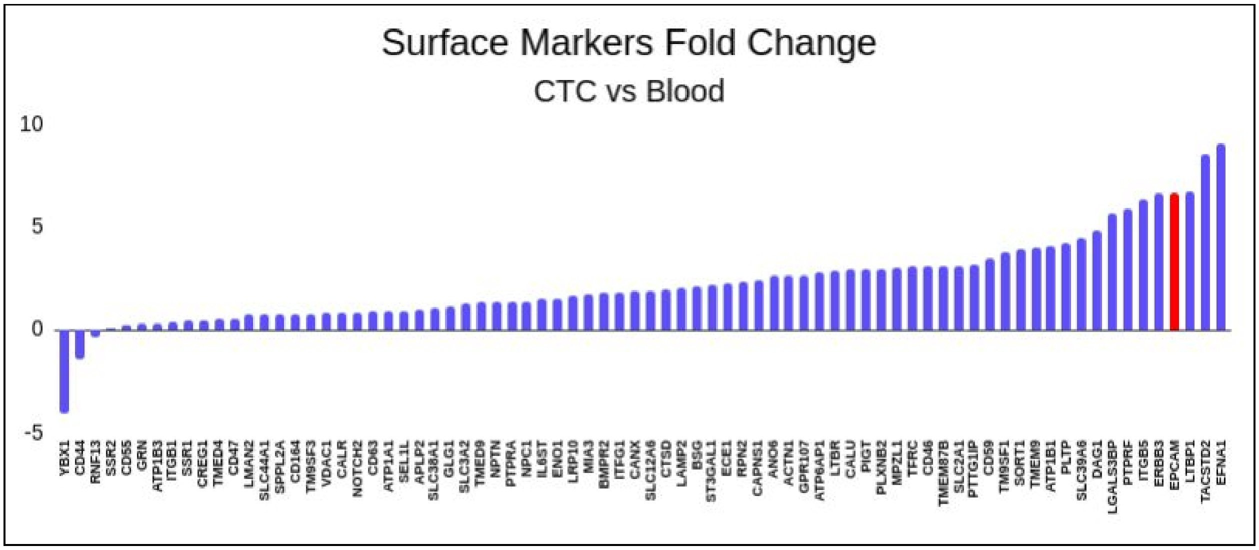


**Supplementary Figure-12:** Log2 fold change of surface markers between CTC and PBMC populations. Besides EpCAM, few genes including ERBB3, LTBP1, TACSTD2, EFNA1 appear specific to CTCs.

**Supplementary Table 1. List of all studies from which datasets are used**

The table can be found as a separate file **Supplementary_Table_1.xlsx.** The sheet contains data of the studies used in the project in the form of a table with identifier, title, number of samples, link etc. More information about the studies can be fetched from the links provided.

**Supplementary Table 2. Functional details of the EMT related genes used in the study.**

The table can be found as a separate file **Supplementary_Table_2.xlsx.** The sheet contains data of the genes used for all the analysis related to EMT.

**Supplementary Table 3. Lists of hybrid E/M gene pairs implicated in cancer survival**

The table can be found as a separate file **Supplementary_Table_3.xlsx.** The sheet contains data of the details of hybrid E/M gene pairs along with their p-values affecting cancer survival. The workbook contains different four different sheets for each cancer.

**Supplementary Table 4. Genes used as features for the machine learning based analyses.** The table can be found as a separate file **Supplementary_Table_4.xlsx.** The sheet contains data of the list of genes used as features for machine learning model.

**Supplementary Table 5. Machine learning results** The table can be found as a separate file **Supplementary_Table_5.xlsx and Supplementary_Table_6.xlsx** The workbook **Supplementary_Table_5.xlsx** contains testing and training statistics done on three ml models for one study out protocol and **Supplementary_Table_6.xlsx** contains testing and training statistics done on three ml models for two studies out method.
